# Deep Sequencing of 10,000 Human Genomes

**DOI:** 10.1101/061663

**Authors:** Amalio Telenti, Levi C.T. Pierce, William H. Biggs, Julia di Iulio, Emily H.M. Wong, Martin M. Fabani, Ewen F. Kirkness, Ahmed Moustafa, Naisha Shah, Chao Xie, Suzanne C. Brewerton, Nadeem Bulsara, Chad Garner, Gary Metzker, Efren Sandoval, Brad A. Perkins, Franz J. Och, Yaron Turpaz, J. Craig Venter

## Abstract

We report on the sequencing of 10,545 human genomes at 30-40x coverage with an emphasis on quality metrics and novel variant and sequence discovery. We find that 84% of an individual human genome can be sequenced confidently. This high confidence region includes 91.5% of exon sequence and 95.2% of known pathogenic variant positions. We present the distribution of over 150 million single nucleotide variants in the coding and non-coding genome. Each newly sequenced genome contributes an average of 8,579 novel variants. In addition, each genome carries in average 0.7 Mb of sequence that is not found in the main build of the hg38 reference genome. The density of this catalog of variation allowed us to construct highresolution profiles that define genomic sites that are highly intolerant of genetic variation. These results indicate that the data generated by deep genome sequencing is of the quality necessary for clinical use.

**Significance statement:** Declining sequencing costs and new large-scale initiatives towards personalized medicine are driving a massive expansion in the number of human genomes being sequenced. Therefore, there is an urgent need to define quality standards for clinical use. This includes deep coverage and sequencing accuracy of an individual’s genome, rather than aggregated coverage of data across a cohort or population. Our work represents the largest effort to date in sequencing human genomes at deep coverage with these new standards. This study identifies over 150 million human variants, a majority of them rare and unknown. Moreover, these data identify sites in the genome that are highly intolerant to variation - possibly essential for life or health. We conclude that high coverage genome sequencing provides accurate detail on human variation for discovery and for clinical applications.

---

Recent technological advances allowed for the large scale sequencing of the whole human genome (1–7). Most studies generated population-based information on human diversity using low to intermediate coverage of the genome (4x to 20x sequencing depth). The highest coverage (30x or greater) was reported for the recent sequencing of 1,070 Japanese subjects (6), 129 trios from the 1000 Genome Project (3), and 909 Icelandic subjects (4). High coverage, also described as deep coverage, may be needed for an adequate representation of the human genome.

In an effort to evaluate the capabilities of whole human genome sequencing using short read sequencing in full production mode, we first measured accuracy and generated quality standards by analysis of the reference material NA12878 from the CEPH Utah reference collection *(8)*. We then assessed these quality standards across 10,545 human genomes sequenced to high depth. This generated a reliable representation of human single nucleotide variation (SNV), and the reporting of clinically relevant SNVs. We confronted, like other groups, the limitations of short read sequencing for accurate calling of structural and copy number variation; even with a variety of methods, resolving structural variation in a personal genome remains a challenge (9).

## Results

**Reproducibility of sequencing on a reference genome.** We assessed the extent of genome coverage using data from 325 technical replicates of NA12878 at different depths of read coverage. The canonical NA12878 Genome-In-A-Bottle call set (GiaB v2.19) defines a set of high confidence regions that corresponds to approximately 70% of the total genome. Regions of low complexity (e.g. centromeres, telomeres and repetitive regions) and challenging for sequencing, alignment and variant calling methods are excluded from the GiaB high confidence region. At the target mean coverage of 30x, 95% of one NA12878 genome is covered at least at 10x. In contrast, at a target mean coverage of 7x used by several genome projects, only 23% of the NA12878 genome is sequenced at an effective 10x (Fig. 1*A*).

**Fig. 1.**
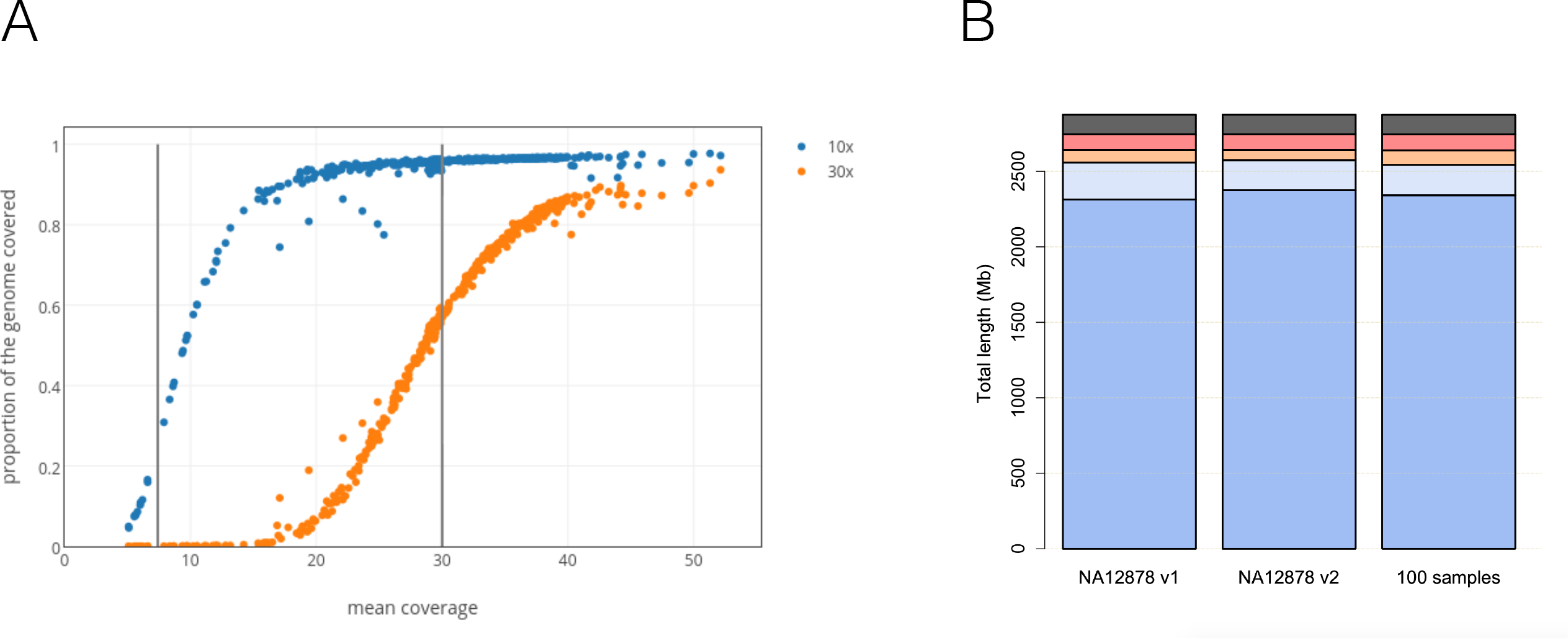
Effective genome coverage and sequence reproducibility. ***(A)***Analysis of the relationship of mean coverage with effective genome coverage uses 100 NA12878 replicates with coverage <30x, 200 replicates with mean coverage of 30x to 40x, and 25 replicates with >40x. Vertical grey lines highlight mean target coverage of 7x and 30x. Each sequencing replica is plotted at lOx (blue) and 3 Ox (orange) effective minimal genome coverage. ***(B)*** Analysis of reproducibility uses NA12878 genomes at 30x-40x mean coverage (two clustering chemistries, vl and v2, each n=100 replicas) to assess the consistency of base calling at each position in the whole genome. The analysis of reproducibility is then extended to 100 unrelated genomes (25 genomes per main ancestry group, African, European, Asian, and for 25 admixed individuals). The color bars represent degree of consistency (blue 100%, light blue ≥90%, orange ≥10-<90%, red <10%, black, noPASS).

We next assessed reproducibility on variant calling for the whole genome by restricting the analysis to a set of 200 samples of NA12878 that were sequenced at a mean coverage of 30x to 40x. Due to manufacturer’s changes in clustering reagents, we analyzed 100 samples prepared with v1 and 100 with v2 (supporting information). After applying quality filters, passing genotypes calls were compared for consistency (Fig. 1*B*). For v2 chemistry, 2.51 billion positions passed, and were called with 100% reproducibility in all replicates. An additional 210 Mb of genome positions yielded passing reproducible genotypes in more than 90%. Only 184 Mb of genome positions were sequenced with lower reproducibility (<90%). Similar results were obtained for v1 chemistry. The analysis of 100 unrelated genomes (25 individuals for each of the three main populations, African, Asian, European, and 25 admixed individuals) confirmed the consistency of calls across genomes (Fig. 1*B*). Overall, a total of 2,157 Mb (97.3%) of the GiaB high confidence region could be sequenced with high reproducibility (Table S1) with a low false discovery rate of 0.0008, precision of 0.999 and recall of 0.994. Overall, these analyses indicate that the current technology and sequencing conditions generate highly accurate sequence data over a large proportion of the genome.

The full extent of sequence generated for a single genome is greater than what is defined by the boundaries of GiaB. It should be noted that the various genome sequencing initiatives use different reporting of what is sequenced (“accessible genome”), what is sequenced confidently, and whether these estimates are reported for an individual genome, or for the collective analysis of multiple genomes. Our work specifically presents the genome calls for a single individual benchmarked against the complete sequence (hg38-Total chromosomal length of autosomes and chrX, 3,031 Mb), and against the community standard (GiaB, on autosomes+chrX, liftover to hg38, 2,215 Mb); Table S2. For a single individual, we map sequence on 90-95% of the genome-and 84% of a single genome is reported at high confidence (see below). In contrast, several published sequencing projects (2–5) describe genome coverage computed from the combination of all genomes-not for an individual genome. Using similar metrics as those in the current work for one 7X mean coverage 1000 Genome Project sample (HG02541), we find the following statistics: 0.2% of the genome covered at 30X, 22.1% covered at 10X, and 75.5% covered at 5X. The loss of coverage genome wide translates in loss of coverage of particular genes and variants. For example, the American College of Medical Genetics and Genomics recommends that laboratories performing clinical sequencing seek and report mutations of 56 genes (10). At 7X mean coverage none of the exonic bases of HG02541 could be covered at 30X, 30% would be covered at 10X and 84% at 5X. Therefore, low coverage genomes are not suitable for clinical use because they can only generate confidence sequence for a fraction of the genome.

**The metrics of 10,000 genomes.** The confidence regions established from sequencing of NA12878 and for 100 unrelated genomes served to guide the analysis of 10,545 human genomes. These samples cover various human populations, admixture and ancestries (Fig S1). We first defined an *extended confidence region* (ECR) that includes the high confidence GiaB regions and the highly reproducible regions extending beyond the boundaries of GiaB (Fig S2). The ECR encompasses 84% of the human genome, and it includes 91.5% of the human exome sequence (Gencode, 96 Mb), which is consistent with recent reports on coverage of the human exome in whole genome analyses *(11)*. We also examined the relevance for clinical variant calls: 28,831 of 30,288 (95.2%) unique ClinVar and HGMD pathogenic variant positions are found in the ECR. We have now confirmed that 373 Mb (86%) of the additional 435 Mb of confident sequence in the ECR is also defined as high confidence in the soon to be released GiaB v3.2.

For 10,545 genomes, the ECR included over 150 million SNVs at 146 million unique chromosomal positions The mean SNV density in the ECR is 56.59 per 1 kb of sequence. However, there are differences across chromosomes: Chr.1 is the least variable (55.12 SNVs/kb) and Chr.16 the most variable (61.26 SNVs/kb) of the autosomal chromosomes. SNV density on ChrX is 35.60 SNVs/kb, but this estimate only considers female genomes (N=6320). A lower mutation rate of variation on the X chromosome than on autosomes is thought to reflect purifying selection of deleterious recessive mutations on hemizygous chromosomes (12). Diversity is further reduced by the effective population size of the X chromosome, because males only carry one copy (13). The SNV density on Chr.Y is 12.70 SNVs/kb, also consistent with knowledge *(14);* however, only male genomes (N=4,225) are considered here and only 15% of the single Y chromosome is included in the ECR (Fig. S3). The definition of ECR allowed for more high confidence calls than those identified in GiaB (Table 1). This is illustrated by the confident identification of 3,390 ClinVar and HGMD pathogenic variant sites identified in the 10,545 genomes: 2,628 (77.5%) were called in the GiaB region, while 3,191 (94.1%) could be called in the ECR (Table 1).

**Patterns of genetic variation in the coding and non-coding genome.** The volume of data presented here provides careful detail on the pattern of sequence conservation and SNVs across the human genome. We compared the rates of diversity in protein coding, RNA coding and regulatory elements (Fig. 2*A*, Fig. S4 and Table S2). On average, protein coding elements are more conserved than intergenic regions and, as previously reported, alternative exons are the least variable (15). Alternative introns of lncRNAs are the most conserved and snoRNA the most variable of RNA coding elements. Among the analyzed DNA regulatory elements, repressed chromatin are the most conserved, and promoters are the least conserved (Fig. 2*A*). There is an extensive literature on the uneven distribution of SNV density across the genome. Positive selection, nucleotide composition, recombination hot spots and replication timing are considered to be contributing factors (16–18). More recently, the sequence context has been shown to explain >81% of variability in substitution probabilities *(19)*. These considerations notwithstanding, the pattern of SNV density is relatively stable across chromosomes (Fig. S5). However, we identified three unique hypervariable megabase-long regions on autosomes (Fig. S6). We observe the depletion of enhancer associated histone marks (H3K4me1, H3K4me2, H3K4me3, H3K27me3 and H3K27ac) in these regions. The hypervariable regions are also gene-poor, and depleted in chromatin loops, leading us to infer that these are domains that are not involved in long-distance interactions between regulatory elements and target genes. The enrichment of variation suggests there is limited purifying selection compared to other regions in the genome.

**Fig. 2.**
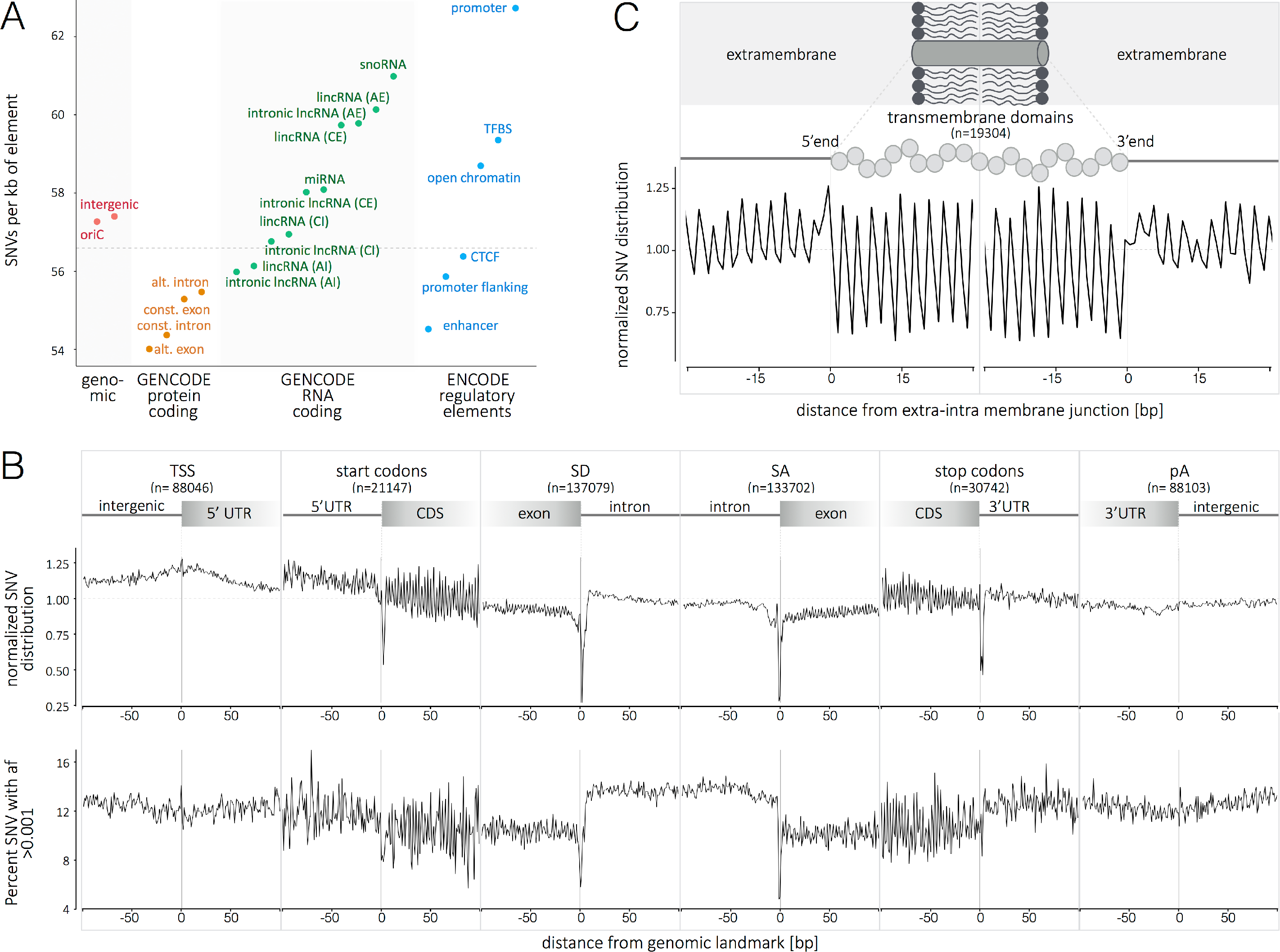
Single nucleotide variant distribution and metaprofiles in the coding and non-coding genome. ***(A)*** Distribution of SNVs in selected genomic elements (genomic, protein coding, RNA coding and regulatory elements; see supporting information for details). The genome average of 56.59 SNVs per kb is indicated by the horizontal dashed line. ***(B)*** The metaprofiles of protein coding genes are created by aligning all elements of 6 different genomic landmarks (TSS, start codon, SD, SA, stop codon and pA) for all 10,545 genomes. The y axis in the upper representation describes the enrichment/depletion of SNVs occurrence per position, normalized to the mean of the protein-coding score (indicated by the horizontal dashed line); the y axis in the lower representation describes the percent of SNVs at each position with an allelic frequency higher than 1 in a 1000. The x axis represents the distance from the genomic landmark. The vertical line indicates the genomic landmark position. The SD and SA metaprofiles highlight the strong conservation of the splice sites (upper panel) and the difference in SNV allele frequency between exons and introns (lower panel). ***(C)*** The metaprofile of Transmembrane domains is created by aligning all single domains at their 5’-and 3’-ends. The figure highlights that every aminoacid in the transmembrane domain is conserved compared to the surrounding structure of the protein. ***(D)*** The metaprofiles of TFBS are created by aligning all the binding sites of four transcription factors (FOXA1, STAT3, NFKB1, MAFF) for all 10,545 genomes. The x axis represents the distance from the 5’ end of the TFBS. The vertical lines indicate the 5’ and 3’ ends of the TFBS. ***(E)*** Ranking of 39 TFBS by conservation (minimum score for the motif; i.e., the nucleotide with lowest tolerance to variation). For panels C-E, the *y* axis describes the normalized enrichment/depletion of SNVs occurrence per position, normalized to the mean of the protein-coding score (indicated by the horizontal dashed line). AE, alternative exon; AI, alternative intron; CE, constitutive exon; CI, constitutive intron; oriC, origin of replication; TSS, transcription start site; SD, splice donor site; SA, splice acceptor site; pA, poly adenylation site; TFBS, transcription factor binding site.

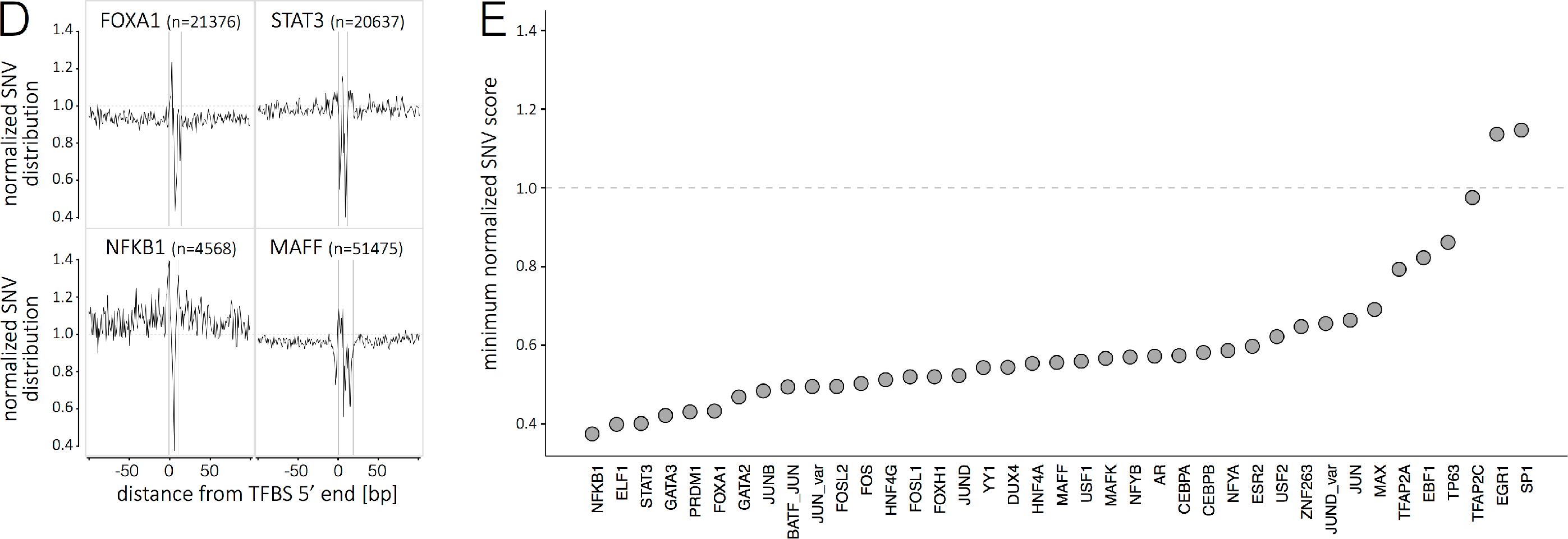

In order to explore the pattern of variation in the human genome in depth, we built “SNV metaprofiles” by collapsing all members of a family of genomic elements into a single alignment. Metaprofiles of protein-coding genes used GENCODE annotated TSS (n=88,046), start codons (n=21,147), splice donor and acceptor sites (n=137,079 and 133,702, respectively), stop codons (n=37,742) and polyadenylation sites (n=88,103) (see supporting information for details). For each nucleotide aligned against these landmark positions, all of the genomes in this dataset (n=10,545) were used to generate a precise representation of the pattern of conservation, and allele spectra (Fig. 2*B*). A pattern is built by incorporating up to 1.4 billion data points (number of aligned elements x 10,545 samples) per genomic position. For example, the analysis captures the decrease in variant allele frequency in exons, with the maximum drop occurring at the splice donor site (Fig. 2 *B*). Positions that do not tolerate human variation can be interpreted as essential and possibly linked to embryonic lethality. In addition, the metaprofiles reveal discreet patterns, including with great precision the periodicity of conservation in coding regions due to the degeneracy of the third nucleotide in the codon in every exon window. The precision of the approach is also illustrated by the metaprofile of 19304 transmembrane domains from 4719 proteins. The constrain of maintaining alpha-helices (or other structures), and the hydrophobic (or polar) nature of the transmembrane domain, result in all aminoacids being distinctively conserved (Fig. 2*C*).

Many differences across individuals and species occur at the level of transcription factor binding *(20)*. We use the binding site core motifs for metaprofile landmarking to identify signatures that include both variation-intolerant and hyper-tolerant positions at the binding site (Fig. 2*D*). Ranking of 39 TFBS by the minimum score of the metaprofile (i.e., the nucleotide with lowest tolerance to variation) emphasizes profound differences in the requirements for conservation across transcription factors (Fig. 2*E*). While the identification of conserved, intolerant sites is expected, the biology behind unique hypertolerant positions at transcription binding sites remains to be investigated.

**Metaprofile tolerance score and variant pathogenicity.** Rare human variants at intolerant sites may carry a greater fitness cost, associate with greater phenotypic consequences and thus can be prioritized for clinical assessment. To apply metaprofiles for scoring of functional severity of variants, we established a tolerance score (Fig. 3*A*) that summarizes the rates and frequency of variation at a given position and for a given landmark. Using this approach, Fig. 3 *B* illustrates the accumulation of pathogenic variant calls at sites with the lowest metaprofile tolerance scores. To formalize this analysis, we calculated the tolerance score at positions aligned to the main coding region landmarks: 100 positions upstream and downstream of the TSS, start codon, splice donor and acceptor, stop codon and polyadenylation site. At the lowest tolerance score, we observe up to 6-fold enrichment for pathogenic variants (Fig. 3*C*). To understand the characteristics of genes that tolerate variants at privileged sites we used an orthogonal assessment of gene essentiality *(21)*. The set of essential genes includes highly conserved genes that have fewer paralogs, and are part of larger protein complexes. Essential genes also display a higher probability of CRISPR-Cas9 editing compromising cell viability (22), and knockouts in the mouse model are associated with increased mortality (23). Fig. 3*D* supports the concept that the less essential genes can tolerate variation at sites with low metaprofile tolerance scores.

**Fig. 3.**
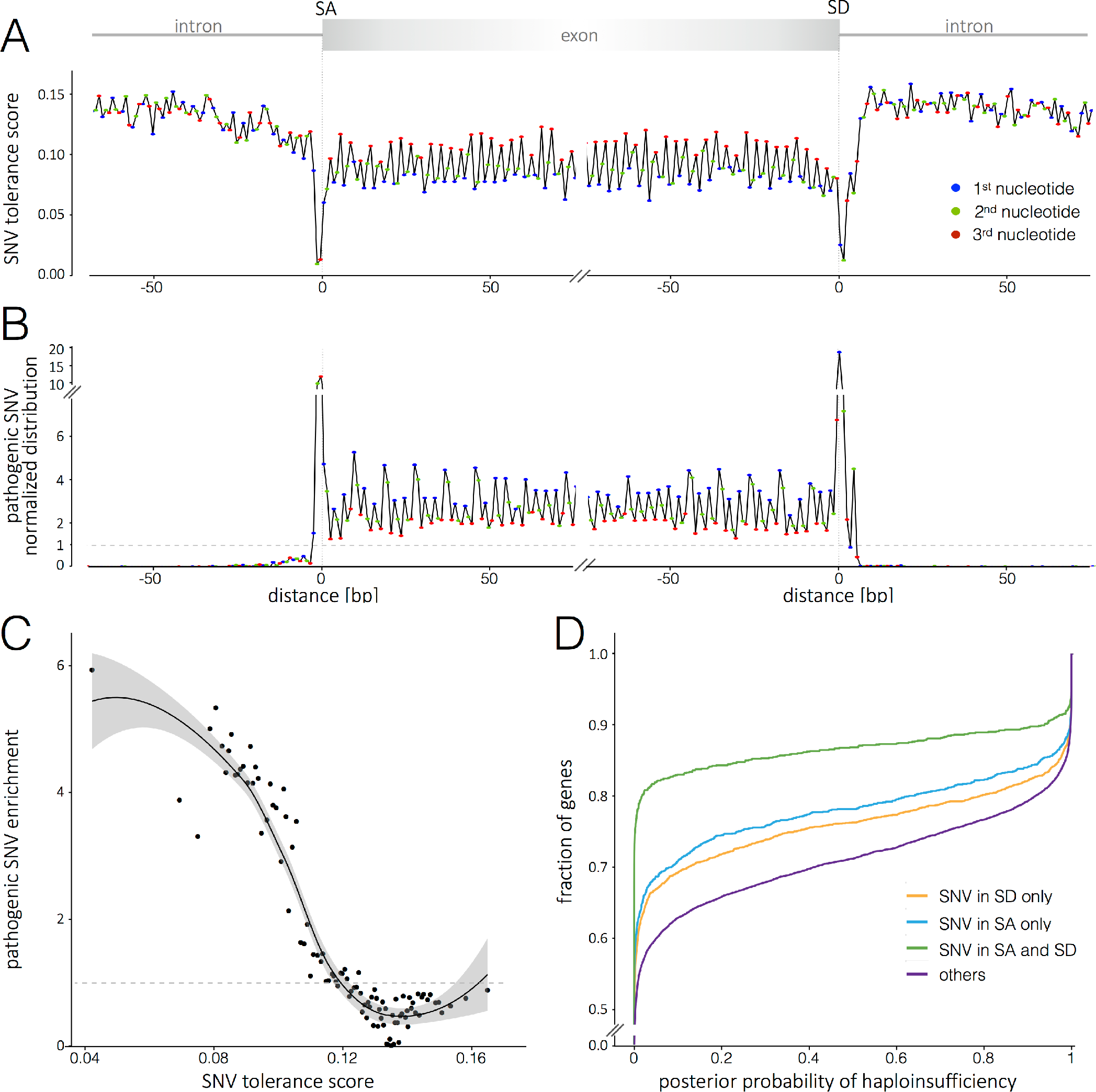
Relationship of a metaprofile tolerance score with variant pathogenicity and gene essentiality. ***(A)***] Metaprofile of the transition between introns and exons expressed as *Tolerance Score* (TS). The TS is the product of the normalized SNV distribution value by the proportion of SNVs with allele frequency ≥ 0.001 (see Fig.2B). The exon sequence highlights the conservation and tolerance to variation of the third position in codons (red). The pattern of higher tolerance to variation every third nucleotide is lost in introns. The TS is lowest at the splice donor and acceptor sites and highest in introns. ***(B)*** The distribution of ClinVar and HGMD pathogenic SNVs (n=29,808 in SD; n=3 0,3 69 in SA metaprofiles) reflects a significant enrichment of pathogenic variants at the sites of lowest TA. Consistently, the exon sequence highlights the enrichment for variation at the first position in codons (blue), as it results in amino acid change or truncation. ***(C)*** Relationship of tolerance score and enrichment for pathogenic variants. Represented on X axis are the median TS values of 1200 positions (six protein coding landmark positions ± 100 bp) expressed in 100 bins. The Y axis present the fold enrichment in pathogenic variants per bin. The LOESS curve fitting is represented by the solid line; the shaded area indicates the 95% confidence interval. ***(D)*** Less essential genes tolerate variation at sites with lowest TS values. The x axis represents a gene essentiality score (the posterior probability of intolerance to truncation *(21)*). The y axis represents the fraction of genes with a given essentiality score or lower. Purple = genes with no variation in splice donor (SD) or acceptor (SA) sites are more essential and appear shifted to the right of the plot, Orange = genes with variation only in SD sites, Blue= genes with variation only in SA sites, Green = genes with variation in SD and SA sites are less essential and appear shifted to the left of the plot.

An important feature of metaprofiling is that it predicts functional consequences of variation solely on the basis of human diversity. In contrast, the Combined Annotation-Dependent Depletion (CADD) score *(24)* uses evolutionary information, annotation from Ensembl Variant Effect Predictor, and extensive information from UCSC genome browser tracks. Despite these profoundly different approaches, the tolerant scores obtained from metaprofiles in protein coding regions perform similarly to CADD for the identification of functional variants (Fig. S7). This observation underscores the potential of metaprofiling to analyze the genome with minimal pre-existing knowledge-in particular in the non-coding genome, as metaprofile tolerance scores only rely on human variation.

**Variant discovery rates per individual.** The large number of genomes, and the coverage of various human populations served to describe the rate of newly observed, unshared SNVs for each additional sequenced genome. We restricted the analysis to the 8,096 unrelated individuals among the 10,545 genomes (supporting information). There is an expectation of 500 million variants identified after sequencing the genomes of 100,000 individuals (Fig. 4*A*). This analysis establishes at the whole genome level prior estimates from the study of a limited set of genes or using exome analysis *(25, 26)*.

**Fig. 4.**
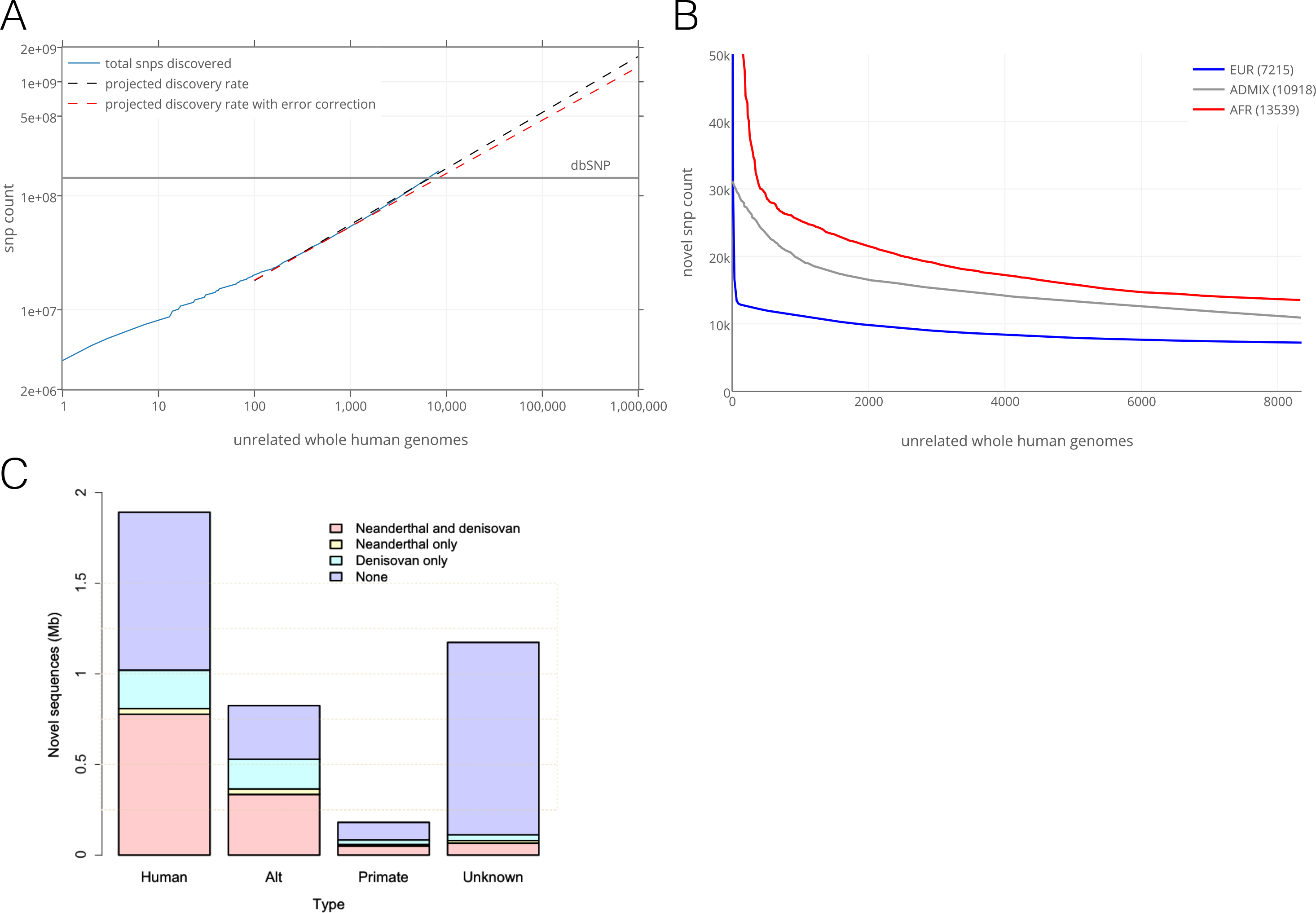
Novel variants and genome sequences. ***(A)*** SNV discovery rate for 8,096 unrelated individual genomes contributing over 150 million SNVs (blue line). The projection for discovery rates as more genomes are sequenced is represented without (dashed black line) and with correction for the empirical false discovery rate of 0.0025 (dashed orange line). The number of SNVs in dbSNP is represented by the horizontal straight grey line. ***(B)*** The number of newly observed variants as more individuals are sequences is determined by the ancestry background and number of participants in the study. Shown are the rates of identification of novel variants for each additional African genome (13,539 SNVs), and for each additional genome of admixed individuals (10,918 SNVs). The most numerous population in the study, Europeans, contribute the lowest number of novel variants (7,215 SNVs). ***(C)*** Unmapped sequence from the analysis of 8,096 unrelated individual genomes contributing over 3.2 Mb of non-reference genome. The 4,876 unique non-reference contigs had matches in NCBI nt database as human, or non-human primate, and with hominins. There are contigs with human-like features that do not have a known match in databases.

Unrelated individuals were assigned to five superpopulations or to an admixed or “other” population group on the basis of genetic ancestry (Fig. S1). Each subsequently sequenced genome contributes on average 8,579 novel variants, that varied from 7,214 in Europeans and 10,978 in admixed, to 13,530 in individuals of African ancestry (Fig. 4*B*). This reflects the current understanding of Africa as the most genetically diverse region in the world *(5)*. Of the 150 million SNVs observed in the ECR, 82 million (54.7%) have not been reported in dbSNP of the National Center for Biotechnology Information or in the most recent phase 3 of the 1000 Genome Project *(3)*. The proportion of novel variants increase with decreasing allele frequency-as expected, there is a negligible number of “novel” variants with allele frequencies greater than 1% (Fig. S8).

**Unmapped human genome sequences.** In addition to new variants, we identified 4,876 unique Human, or human-like contigs (supporting information) assembled from 3.26 Mb of non-reference (hg38 build) sequences (“unmapped reads”). On average, we identified 0.71 Mb of non-reference sequences per genome. A total of 1.89 Mb of the non-redundant sequences could be mapped to known human sequences in GenBank (though not in the hg38 reference assembly). An additional 0.18 Mb mapped to primate sequences in the NCBI nt database. There are 1.17 Mb that did not have a known match in nt or nr databases. The GC content and dinucleotide bias of the unknown contigs reflect the patterns of human sequences. However, we also identified successfully mapped Eukaryotic, Prokaryotic and viral contigs that had indistinguishable metrics from human contigs (Fig. S9). Therefore, it remains difficult to solve bioinformatically the nature of unmapped human-like reads-they may simply result from contamination *(27)*. Much of the non-reference sequence is shared with hominins. The unmapped contigs were compared to Neanderthal and Denisovan sequencing reads that did not map to hg38. There were 0. 96 Mb covered by Neanderthal reads and 1.18 Mb covered by Denisovan reads. In addition, 0.82 Mb are not in hg38 primary assembly, but in the “alt” sequences or subsequent patches (Fig. 4*C*). The presence in some individuals of novel sequence content that is also found among unmapped reads from Denisovan and Neanderthal genomes and in non-human primates reinforce the notion that the human genome is larger and more distributed than what is currently represented by a single (hg38) reference genome.

## Conclusions

The goal of clinical use of the genome requires standards for sequencing, analysis, and interpretation. Our work specifically addresses the first two steps: sequencing and sequence analysis. The performance of the platform, implemented in full production mode, improves on recent benchmarks for the accurate interpretation of next-generation DNA sequencing in the clinical setting *(26, 28, 29)*. This is needed for laboratory standards, for regulatory purposes, and for clinical diagnostics and research. The third step-interpretation-remains a major issue given the many types of genetic evidence that laboratories consider. Initiatives such as ClinVar, and policies and guidelines (10, 30) set standards for clinical interpretation.

This report also extends prior efforts at genome and exome sequencing by detailing the distribution of human variation in the non-coding genome. The amount of data supports the discovery of sites in the genome that are intolerant to variation. The 10,545 genomes provide estimates of the rate of discovery of new SNVs, and complements the human genome by more than 3Mb through the identification of non-reference and of putative human-like sequences. These data anticipate the relentless accumulation of rare variants and the scale of observable mutagenesis of the human genome.

## Materials and Methods

Detailed material and methods are provided in SI Materials and Methods.

## Acknowledgements

Access to the NA12878 replicates and to genome data is being provided as described in SI Materials. We thank Drs. Thomas Caskey and Michael Hicks for useful commentaries.

## Contributions

J.C.V. conceived the study. A.T. led the analyses. L.C.T.L., J. di I., E.H.M.W., E.F.K., A.M., N.S., and C.X., performed the analyses. W.H.B., M.F. and E.S. led the sequencing process. N.B., G.M., M.D.S., S.C.B, C.G., and C.M., built informatic annotation and technology infrastructures, B.A.P., F.J.O, and Y.T. supervised reseach.

